# Influence of Fast-Spiking Prefrontal Neurons on Working Memory Behavior

**DOI:** 10.1101/2023.05.28.542641

**Authors:** Sophia Chung, Rana Mozumder, Sihai Li, Christos Constantinidis

**Affiliations:** Neuroscience Program, Vanderbilt University, Nashville, TN 37235; Department of Biomedical Engineering, Vanderbilt University, Nashville, TN 37235; Department of Neurobiology, The University of Chicago, Chicago, IL 60637; Department of Ophthalmology and Visual Sciences, Vanderbilt University Medical Center, Nashville, TN 37212

## Abstract

Working memory is a limited-capacity system for maintaining and manipulating information for recall. Neurons that generate persistent activity in the primate dorsolateral prefrontal and posterior parietal cortex have been shown to be predictive of behavior in working memory tasks, though subtle differences between them have been observed in how information was represented, in some tasks. The role of different neuron types in each of these areas has not been investigated at depth. We thus compared the activity of neurons classified as fast-spiking, putative interneurons, and regular-spiking, putative pyramidal neurons, recorded from the prefrontal and posterior parietal cortex of male monkeys, to analyze their role in the maintenance of working memory. Our results demonstrate that fast-spiking neurons are active during a range of tasks and generate persistent activity during the delay period over which stimuli need to be maintained in memory. Furthermore, the activity of fast spiking neurons, particularly in the prefrontal cortex, is predictive of the subject’s recall no less than that of regular-spiking neurons, which are exclusively projection neurons in the cortex and thus capable of transmitting signals from the prefrontal cortex into other areas. Our results shed light onto the fundamental neural circuits that determine subjects’ memories and judgments.

## INTRODUCTION

Working memory is the limited-capacity system of maintaining and manipulating information in current thought^1^. It is applicable in spatial, episodic, and verbal tasks^2^ and can be improved with training^3, 4^. Lesion, neuroimaging, and neurophysiological studies have shown the prefrontal cortex (PFC) and the posterior parietal cortex (PPC) to be two key brain regions that give rise to this cognitive ability^5^. Analysis of neurophysiological recordings in non-human primates during visuospatial working memory tasks, such as the oculomotor delayed-response task, revealed that neurons generate persistent activity which represents information regarding the location of visual stimuli, thus providing a neural correlate of working memory^6^. These studies have allowed for the development of theories regarding the neural mechanisms of working memory^7^. The bump attractor model provides mechanistic insights on how persistent activity can be maintained, by virtue of recurrent excitation between neurons, and accounts for behavior in working memory task, tying fluctuations of persistent firing rate to variability of responses^8, 9^.

However, the neural mechanisms through which working memory arises are a continuing debate in the field. Competing theories have questioned whether persistent activity is present across all tasks, suggesting instead, that “activity-silent” models can better account for working memory^10^. Other models have posited that the rhythmic spiking of neurons is the critical neural variable, instead, so that each attractor state is accompanied by bursts of gamma oscillations, resulting from fast, local feedback inhibition^11, 12^. To adjudicate between competing models, it is thus imperative to determine whether proposed neural correlates of working memory can account for behavior across different cognitive tasks, and similarly, whether model predictions hold across tasks.

In a recent study Li et al. (2021) aimed to test whether persistent activity could predict subtle changes in what a subject recalls and how this activity differed in different regions implicated in working memory, namely PFC and PPC. This study used a novel working memory task that dissociated motor preparation from spatial working memory. PFC and PPC neurons were shown to be predictive of behavior. In a second study of interest, Qi et al. (2015) trained monkeys to remember either the first or second of two stimuli presented in sequence. This study, too, revealed that neurons with persistent activity, particularly in the PFC can account for what information the subjects represented in memory.

These studies examined activity pooled from all neurons in these areas, however it is known that pyramidal neurons and interneurons can play distinct role during cortical processing^15, 16^. The bump attractor model predicts that persistent activity is maintained by virtue of structured connections between pyramidal neurons and interneurons and that both populations generate tuned persistent activity that should be predictive of behavior^8^. Different neuron types can be classified from extracellular recordings based on the waveform of their action potentials as fast-spiking (FS) with shorter action potentials and regular-spiking (RS) with longer action potentials^17, 18^. We were therefore motivated to determine whether FS neurons generate persistent activity across tasks and brain areas, and whether their such persistent activity is predictive of behavior, consistent with the bump attractor model.

## MATERIALS AND METHODS

The following methods are summarized from Li et al. (2021) and Qi et al. (2015). Four male rhesus monkeys (*Macaca mulatta*) were used in these experiments. Neural recordings were carried out in areas 8 and 46 of the dorsolateral prefrontal cortex and areas 7a and lateral intraparietal area (LIP) of the posterior parietal cortex. All experimental procedures followed guidelines by the U.S. Public Health Service Policy on Humane Care and Use of Laboratory Animals and the National Research Council’s Guide for the Care and Use of Laboratory Animals were reviewed and approved by the Wake Forest University Institutional Animal Care and Use Committee.

### Experiment setup

Monkeys sat in a primate chair with their head fixed while viewing a liquid crystal display monitor. Animals fixated on a white square in the center of the monitor screen. Animals were required to fixate on a 0.2° spot appearing in the center of the monitor screen. During each trial, the animals maintained fixation on the spot while visual stimuli were presented at peripheral locations. Any break of fixation terminated the trial, and no reward was given. Eye position was monitored throughout the trial using a non-invasive, infrared eye position scanning system (model RK-716; ISCAN, Burlington, MA). Eye position was sampled at 240 Hz, digitized, and recorded. Visual stimuli display, monitoring of eye position, and the synchronization of stimuli with neurophysiological data were performed with in-house software^19^ implemented in MATLAB (Mathworks, Natick, MA).

### Behavioral Tasks

Two monkeys were trained to perform the Match-Stay Nonmatch-Go (MSNG) task (Fig.1A). The task required the monkeys to remember the location of a cue. After a 3s delay period, a second stimulus appeared, either at the identical location (match) or a different location (nonmatch). After 500 ms, the fixation point changed color. If the second stimulus was a match, the monkey was required to maintain fixation; if the second stimulus was a nonmatch, the monkey was required to make a saccade towards this visible stimulus. The monkeys received a liquid reward for each correct response. Possible cue locations included a reference location (white square in the inset of Fig. 1A) and eight locations deviating from the reference location by an angular distance of 11.25°, 22.5°, 45° and 90°, clockwise and counterclockwise. In each daily session, the cue could appear pseudo-randomly at one of the nine possible locations. Since only a limited range of stimulus locations was explored in each session, an effort was made to position stimuli based on the estimated best neuronal responding location of neurons isolated in real-time, however we recorded neuronal activity with multiple electrode arrays, and the location of the stimuli could fall at any position relative to a neuron’s receptive field. The cue was followed by a matching stimulus appearing at the same location as the cue in approximately half the trials or by a nonmatch stimulus, which could only appear at the reference location. The reference location varied from session to session.

**Figure 1.**
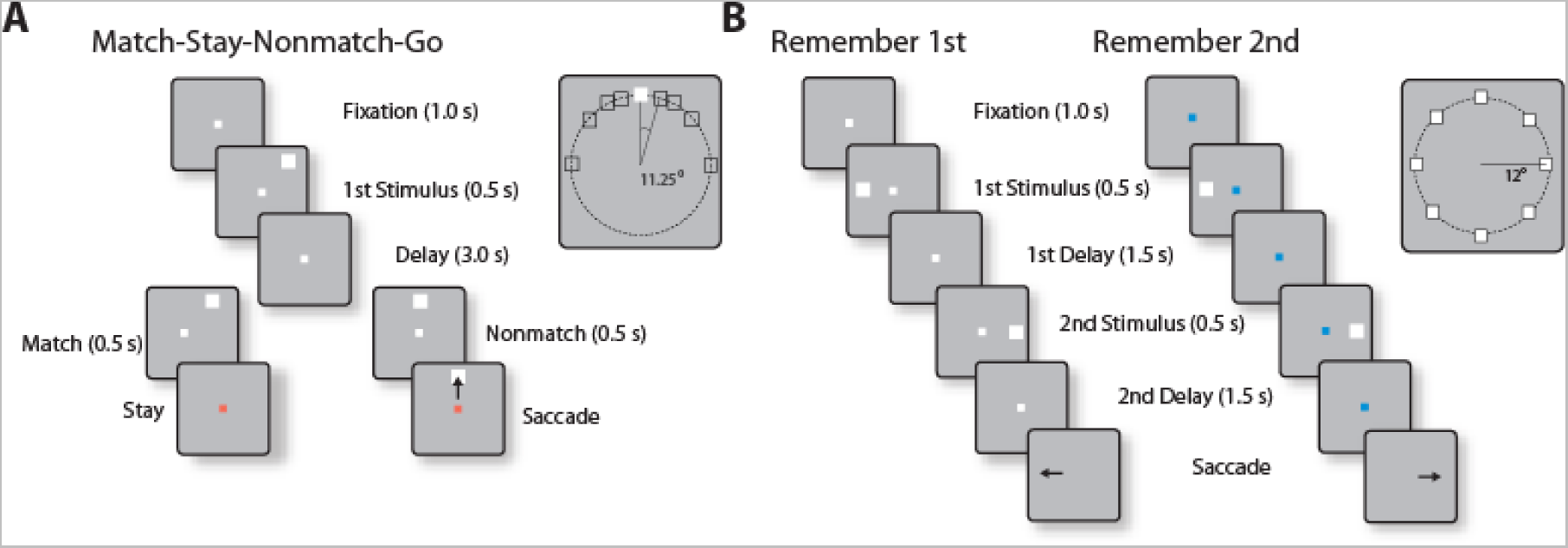
(A) Sequence of events in the MSNG task. The monkey is required to observe the first cue and maintain fixation during the delay period. Then another visual stimulus appears, and the monkey needs to determine if it appeared at the same location as the cue. If it appeared at the same location (match), the monkey needs to stay at the fixation point after the color changes. If the location of the second stimulus deviates from the first (nonmatch), the monkey is required to make an eye movement to the (visible) second stimulus once the color of the fixation point changes. (B) Sequence of events in the Remember First – Remember Second task. In the remember-first task, the fixation point is white, and the monkey is required to make an eye movement to the first stimulus, regardless of the location of a second stimulus, which is a distractor. In the remember-second task, the first stimulus is now a distractor and the monkey is required to make an eye movement to the location of the second stimulus.

Two different monkeys were trained to perform the Remember 1st – Remember 2nd task (Fig. 1B). In this task, two stimuli also appeared in sequence, with intervening delay periods between them, now requiring the monkey to remember and make an eye movement to either the first or the second stimulus according to the color of the fixation point. The monkeys were required to saccade to the location of the first stimulus if the fixation point was white in color (remember-first condition), and to the location of the second stimulus if the fixation point was blue (remember-second condition). To minimize confusion about the stimulus to be remembered, trials with white and blue fixation points were presented in blocks.

### Surgery and neurophysiology

Two, 20-mm diameter craniotomies were performed over the lateral prefrontal cortex and the posterior parietal cortex, and a recording cylinder was implanted over each site.

Neurophysiological recordings were obtained as described before^20^. Tungsten-coated electrodes with a 200 or 250 µm diameter and 4 MΩ impedance at 1 kHz were used (FHC, Bowdoinham, ME). Arrays of up to 4-microelectrodes spaced 0.5-1 mm apart were advanced into the cortex with a Microdrive system (EPS drive, Alpha-Omega Engineering) through the dura into the cortex.

### Neural data analysis

Data analysis was performed in the MATLAB computational environment (Mathworks, Natick, MA, version 2019-2022a). Recorded spike waveforms were sorted into separate units using a semi-automated cluster analysis process of the KlustaKwik algorithm^21^. Neurons were classified as fast-spiking (FS) putative interneurons or regular-spiking (RS) putative pyramidal neurons based on spike width, replicating the method described by Qi et al. (2011), which adapted an earlier method by Constantinidis & Goldman-Rakic (2002). Neurons were classified as FS if they exhibited a spike width of ≤ 550 μs, while those with a spike width ≥ 575 μs were classified as RS.

Neurons generating persistent activity were identified as those with firing rates during the (first) delay period that were higher compared to the 1s baseline fixation period that preceded the cue presentation, based on a paired t-test, evaluated at the p < 0.05 level. Population discharge rates were evaluated by averaging activity from multiple neurons and constructing Peri-Stimulus Time Histograms (PSTH). These were constructed using the best stimulus (preferred cue) for each neuron. Correct and error conditions were compared for trials also involving the preferred cue of each neuron.

To quantify the trial-to-trial association between perceptual choice and neuronal activity, we analyzed trials that resulted in correct choices and incorrect choices using Receiver Operating Characteristic (ROC) analysis^22, 23^. Firing rates of trials involving the same sequences of stimuli were pooled separately for correct and error outcomes. An ROC curve was computed from these two distributions of firing rates. The area under the ROC curve is referred to in the perceptual inference literature as “choice probability” and represents a measure of correlation between the behavioral choice and neuronal activity. A value of 1 indicates a perfect correlation between the behavioral choices and the neuronal discharge rates; a value of 0.5 indicates no correlation between the two. Time-resolved choice probabilities were computed from the spikes in 1000 ms time windows, stepped by 50 ms intervals. Results from all available neurons were averaged together to produce population responses.

## RESULTS

### Persistent activity of FS and RS neurons in the PFC and PPC

Neuronal activity was recorded from areas 8 and 46 of the dorsolateral prefrontal cortex and areas 7a and LIP of the posterior parietal cortex (Fig. 2) in four monkeys. Two of the monkeys were trained to perform the MSNG task (Fig. 1A). A total of 709 neurons were recorded from the PFC and 1017 neurons were recorded from the PFC in this task. Of those, 32 PFC neurons were classified as FS (5%) and 107 PPC neurons were classified as FS (11%). We focused particularly on neurons that exhibited significantly elevated responses in the first cue period or the delay period in the MSNG task, compared with the baseline activity (paired t-test, P < 0.05) as these were identified in Li et al. (2021). A total of 144 neurons with persistent activity following the (limited range of) stimuli used in each session were identified in PFC (49 in subject KE, and 95 in subject LE) and 145 neurons in the PPC (44 in subject KE, and 101 in subject LE). Of those, 11 PFC neurons were identified as FS (8%); similarly, 14 PPC neurons were identified as FS (10%). The proportion of FS neurons that exhibited persistent activity did not deviate significantly from the overall proportion of neurons that were classified as FS in the entire sample for either PFC or PPC (χ^2^-test, p=0.07 and p=0.73, respectively).

**Figure 2.**
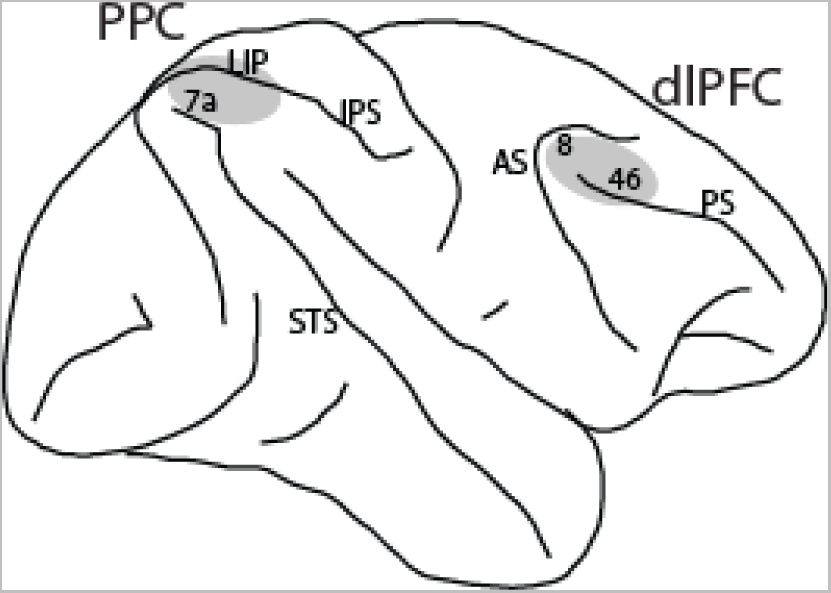
Regions of neurophysiological recordings, comprising areas 8 and 46 in the dorsolateral prefrontal cortex (dlPFC) and areas 7a and LIP in the posterior parietal cortex (PPC). Abbreviations: IPS, intraparietal sulcus; STS, superior temporal sulcus; AS, arcuate sulcus; PS, principal sulcus.

Two additional monkeys were trained to perform the Remember 1st–Remember 2nd task (Fig. 1B). A total of 421 neurons were recorded from the PFC and 551 neurons were recorded from the PPC in this task. Of those, 46 PFC neurons were classified as FS (11%) and 73 PPC neurons were classified as FS (13%). We again determined the neurons that exhibited significantly elevated responses in the first delay period compared with the baseline activity (paired t-test, P < 0.05) as these were identified in Qi et al. (2015). A total of 182 neurons with persistent activity were identified in the PFC (42 in subject GR, and 140 in subject HE) and 179 neurons in the PPC (86 in subject GR, and 93 in subject HE). Of those 15 PFC neurons were identified in FS (8%); similarly, 24 PPC neurons were identified as FS (13%). The proportion of FS neurons that exhibited persistent activity in this task too did not deviate significantly from the overall proportion of neurons that were classified as FS, in either PFC or PPC (χ^2^-test, p=0.24 and p=0.95, respectively). These results suggest that comparable percentages of FS and RS neurons exhibited persistent activity in the context of these working memory tasks, in both the PFC and PPC.

The FS and RS neurons with persistent activity in the two tasks formed the bases of further analysis. The distribution of spike widths from each brain region and experiment is shown in Fig. 3. Although classification into FS and RS categories was based on spike width alone, FS neurons exhibited, on average, higher peak firing rates in both the MSNG task (Fig. 4, 5) and the Remember 1st – Remember 2nd task (Fig. 6). FS neurons generated robust responses to cue stimuli (mean rate for the best cue response of each neuron is shown in Fig. 4) and during the delay period (mean rate for the best delay period response of each neuron is shown in Fig. 5).

**Figure 3.**
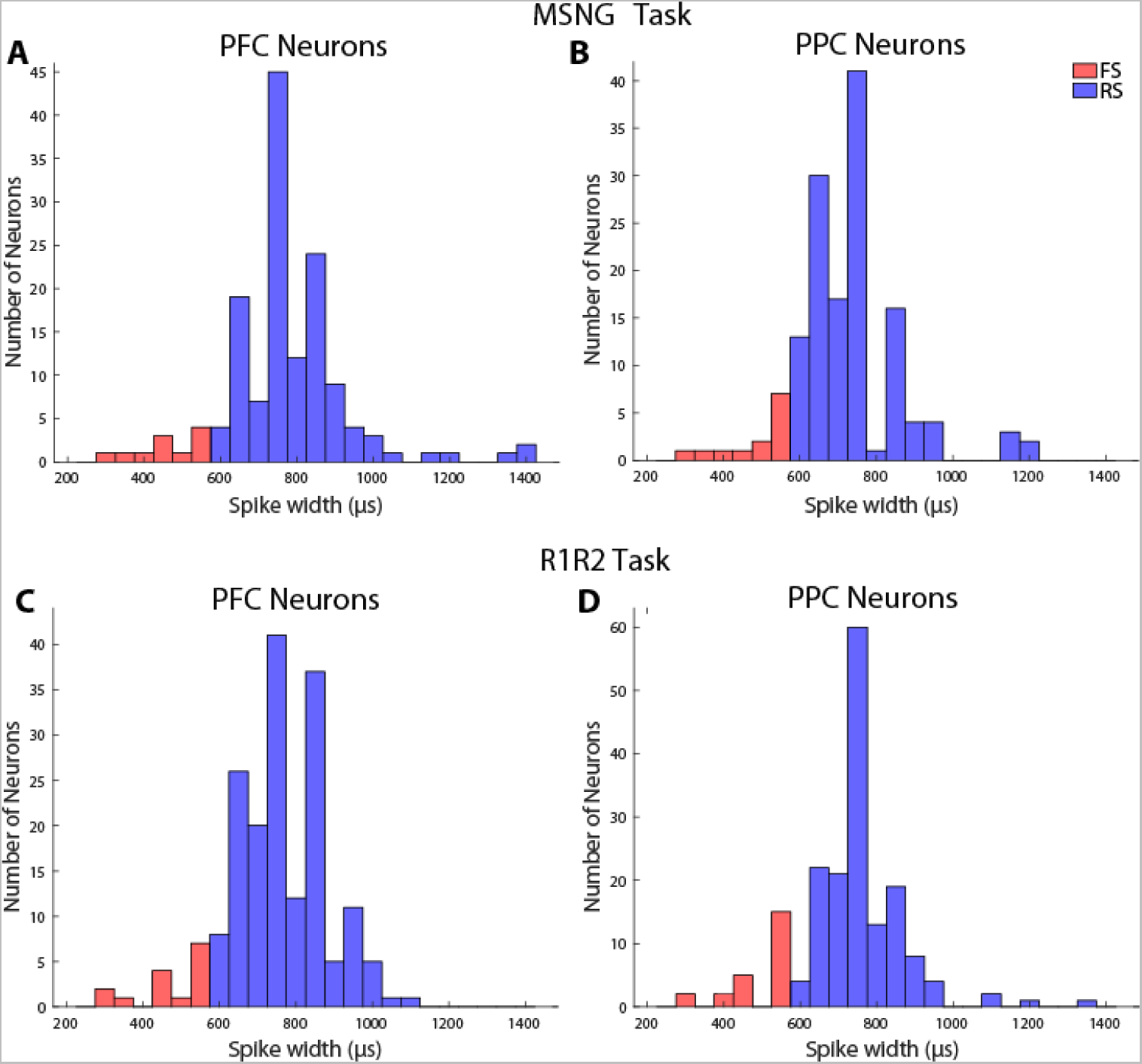
Distribution of spike widths among neurons with significant persistent activity in the delay period of the task. Neurons were classified as fast-spiking (FS) or regular-spiking (RS) according to spike width. (A) Sample size of 11 FS neurons, 133 RS neurons (PFC). (B) 14 FS neurons, 131 RS (PPC) neurons for the MSNG task. (C) Sample size of 15 FS neurons, 167 RS neurons (PFC). (D) 24 FS neurons, 155 RS neurons (PPC) for the Remember First – Remember Second task.

**Figure 4.**
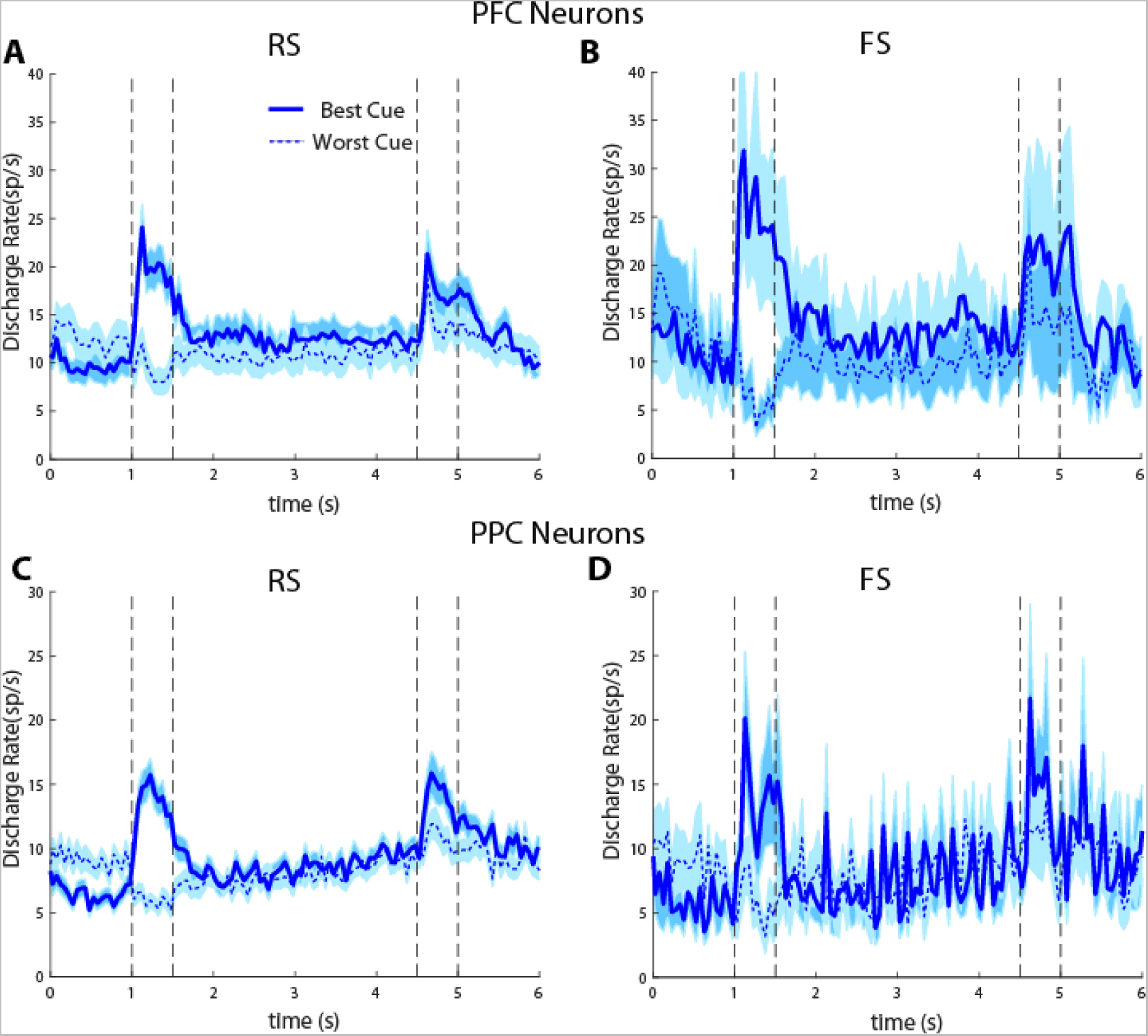
(A) Averaged PSTH of neuronal spike discharges from regular-spiking PFC neurons recorded in the MSNG task (n = 133). The solid lines indicate average activity of RS neurons when the stimulus appeared in their preferred location, while the dotted lines indicate neuronal activity when the stimulus appeared at their worst location tested. Shaded areas indicates standard error of the mean (SEM). (B) Same as in A, for fast-spiking PFC neurons (n = 11). (C) Averaged PSTH of neuronal spike discharges from regular-spiking PPC neurons recorded in the MSNG task (n = 131). (D) Same as in C, for fast-spiking PPC neurons (n = 14).

**Figure 5.**
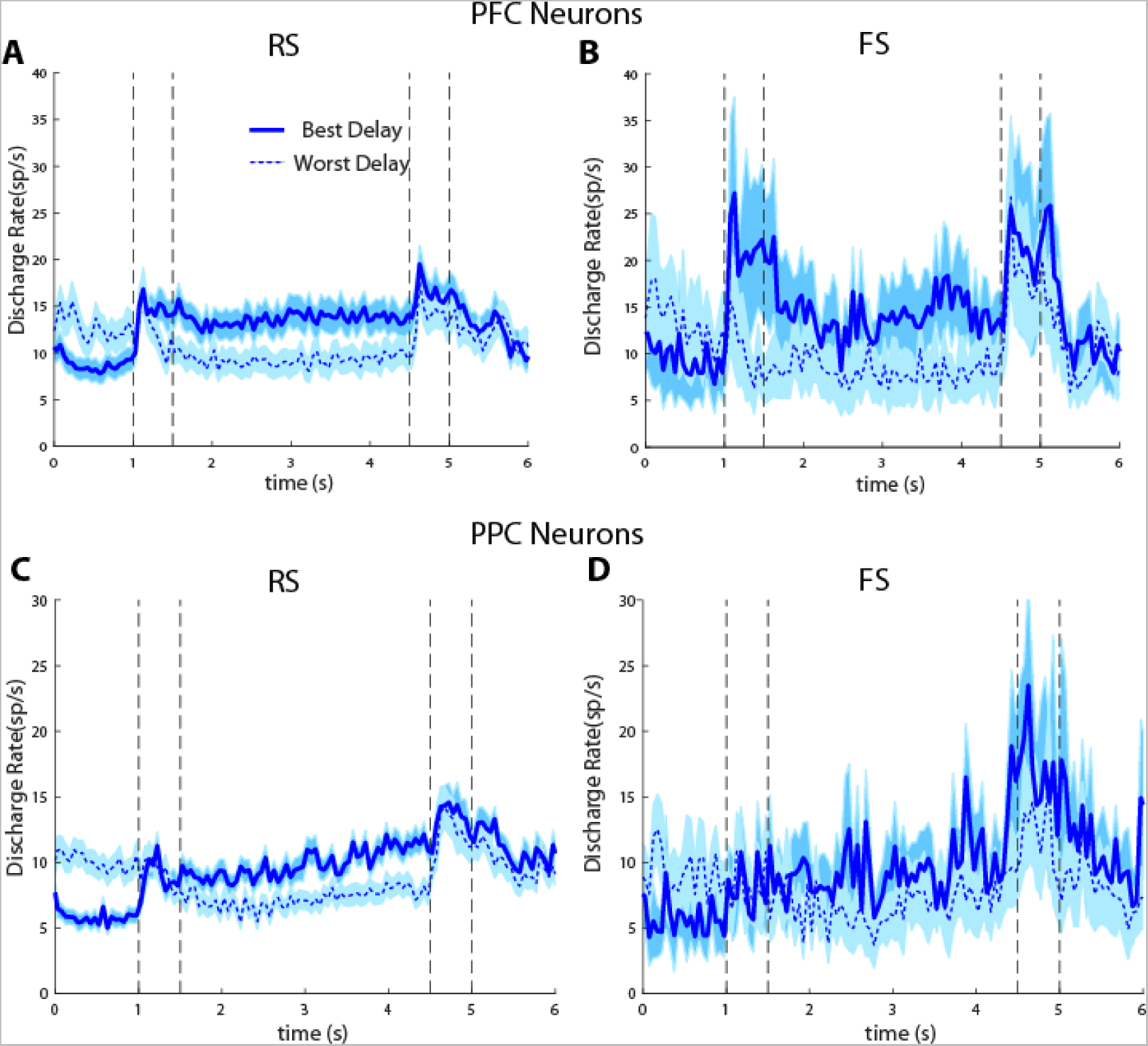
Averaged PSTH of neuronal spike discharges from regular-spiking PFC neurons recorded in the MSNG task (n = 133). Solid lines indicate the neuronal activity of neurons when the stimulus appeared at the location that generated the best delay-period activity, while the dotted lines indicate neuronal activity when the stimulus appeared at the location that generated the weakest delay period activity. Shaded areas indicate the standard error of mean (SEM). (B) Same as in A, for fast-spiking PFC neurons (n = 11). (C) Averaged PSTH of neuronal spike discharges from regular-spiking PPC neurons recorded in the MSNG task (n = 131). (D) Same as in C, for fast-spiking PPC neurons (n = 14).

**Figure 6.**
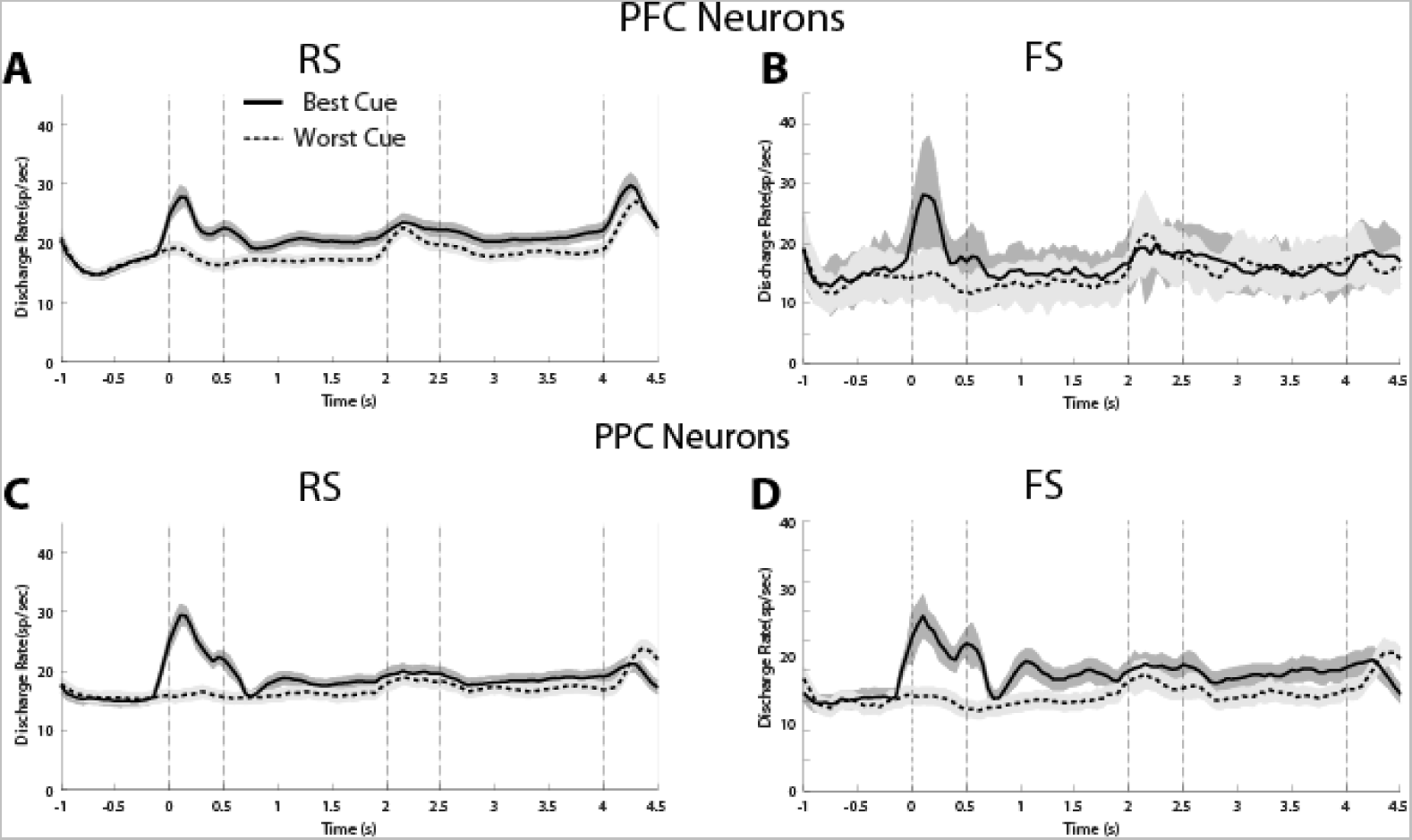
(A) Averaged PSTH of neuronal spike discharges from regular-spiking PFC neurons recorded in the remember first-remember second task (n = 167). Solid lines indicate neuronal activity when the cue appeared at the location that generated the best activity, while the dotted lines indicate activity when the cue appeared at the worst location tested. Shaded areas indicate standard error of mean (SEM). (B) Same as in A, for fast-spiking PFC neurons (n = 15). (C) Averaged PSTH of neuronal spike discharges from regular-spiking PPC neurons recorded in the MSNG task (n = 155). (D) Same as in C, for fast-spiking PPC neurons (n = 24).

### Choice Probability Analysis

To assess the reliability with which firing rates can predict the subject’s choice in the task, we performed a Receiver Operating Characteristic (ROC) analysis that compared the distributions of firing rates in correct and error trials, yielding a quantity sometimes referred to as choice probability^22^. This analysis was performed using data from the preferred cue location of each neuron, averaging choice-probability values across neurons in a time-resolved fashion (Fig. 7). In the MSNG task, which requires memory of a single stimulus and then comparison with a second stimulus we have previously shown that the activity of both PFC and PPC neurons influences the judgments that monkeys make in this task^13^, with very similar properties between the two areas. We therefore began our analysis by pooling FS neurons from both areas and comparing their properties with RS neurons. The overall time course of choice probability was qualitatively similar for the FS and RS neurons. The time interval of maximal choice probability for RS neurons occurred during the second stimulus presentation (arrow in Fig. 7A-B). In other words, if the activity of RS neurons was lower than average in a trial at this time interval, the subject was much more likely to make an erroneous than correct choice. This choice probability value was significantly higher than the chance value of 0.5 (two-tailed, one-sample t-test, t_151_=4.08, p=7.3x10^-5^). We then repeated the analysis for the same time interval for FS neurons. Indeed, for FS neurons, too, choice probability was significantly higher than 0.5 (two-tailed, one-sample t-test, t_12_=2.77, p=0.017). Testing separately PFC and PPC neurons suggested that RS neurons in both the PFC and PPC reached a significant choice probability value during the second stimulus presentation (t_77_=3.0726, p=0.0029 for PFC, t_73_=3.0742, p=0.0030 for PPC). For FS neurons, our sample was much smaller, nonetheless PFC neurons tested alone reached statistical significance (t_6_=3.176, p=0.019) whereas PPC neurons did not (t_5_=1.3, p=0.2).

**Figure 7.**
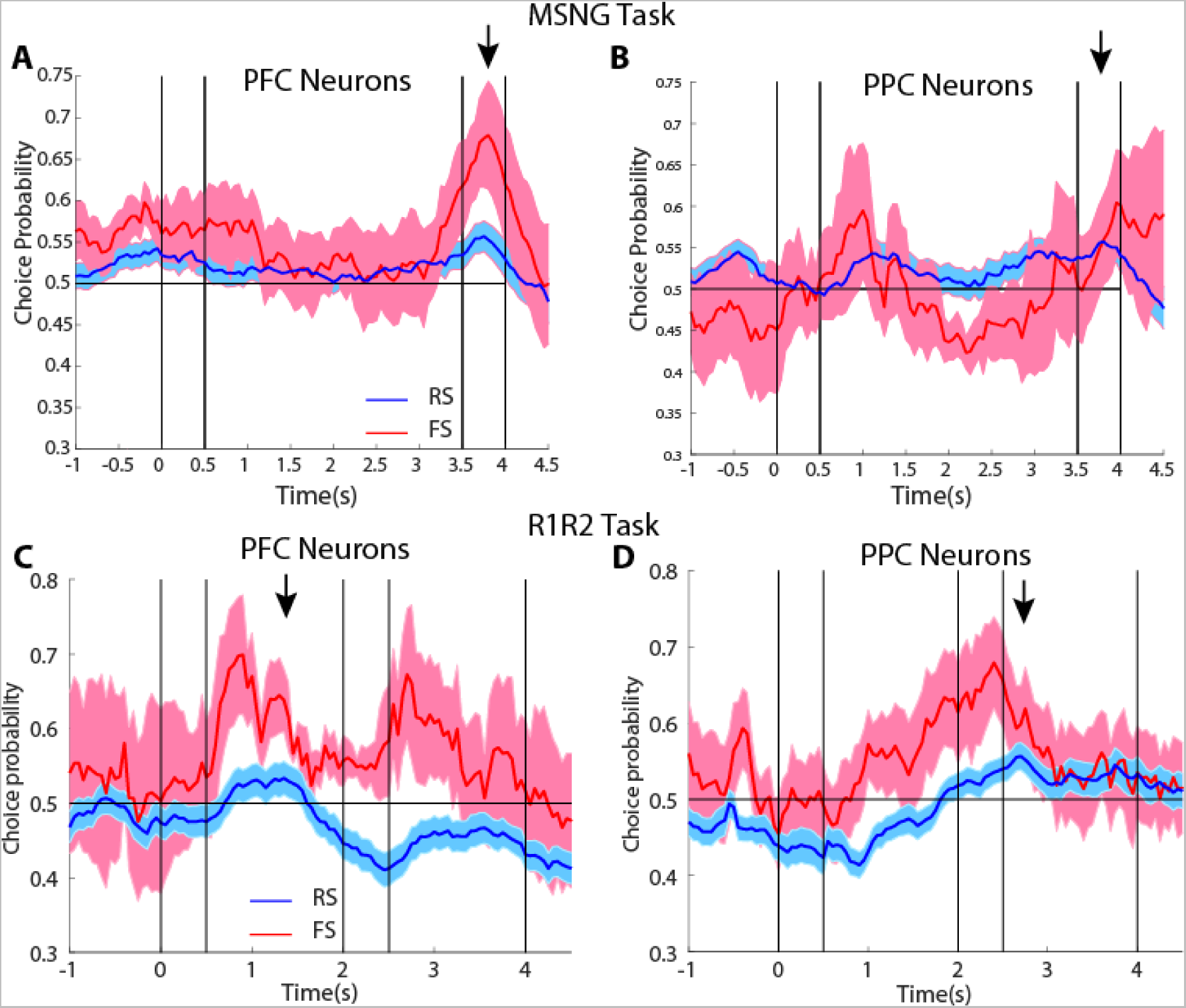
Averaged area under the ROC curve from FS neurons recorded from the PFC (A), and the PPC (B) plotted as a function of time across the trial. The solid line represents ROC value comparing the distribution of correct and error nonmatch trials from the preferred cue condition, the dotted line representing the nonpreferred cue condition. The shaded area around represents SEM.

We repeated this analysis for the Remember First – Remember Second task. In this task, which involves a distractor stimulus that the monkey has to ignore, we have reported that the activity of PFC neurons is much more predictive of behavior^14^. PPC neurons are generally not able to resist the appearance of a distractor, and their responses in this interval were less likely to determine the eventual choice of the subject. In this case, too, the overall time course of choice probability was qualitatively similar for the FS and RS neurons. We determined the time point of maximum choice probability across all PFC neurons in the Remember First condition, which occurred at the beginning of the first delay period (arrow in Fig. 7C). For FS neurons, choice probability was significantly higher than chance at this time interval (two-tailed, one-sample t- test, t_3_=3.27, p=0.047). For PPC neurons, the maximum choice probability was observed at the beginning of the second delay period. RS neurons in that interval exhibited a significantly higher than chance choice probability (two-tailed one-sample t-test, t_70_=3.17, p=0.002). When we examined the FS neurons of the PPC, although a trend in the same direction was evident, this did not reach statistical significance (two-tailed, one-sample t-test, t_12_=1.96, p=0.07).

## DISCUSSION

The persistent activity model is considered the neural correlate of working memory^24^, but continues to be debated^10, 11^. The bump-attractor computational model suggests that persistent activity is sustained by virtue of recurrent connections between neurons with similar tuning, thus allowing discharges to reverberate in the circuit and be maintained even after the actual stimulus is no longer present^8, 25, 26^. Structured excitatory and inhibitory connections are both essential in the maintenance of working memory in this model^27, 28^. Our current results confirm this model prediction and demonstrate that fast spiking, putative interneurons are active during a variety of cognitive tasks and generate persistent activity following stimuli that are required to be remembered.

Persistent activity has been shown to predict working memory performance, however evidence linking behavior with persistent activity based on delayed response tasks has been criticized as it may represent motor preparation rather than working memory per se^29, 30^.

Alternative tasks have been introduced, such as the MSNG task, which requires a categorical judgment about two stimuli and decouples the stimulus that needs to be maintained in memory from the eventual response. Neuronal firing rate deviations during the delay interval of the task have been found to be predictive of what a subject recalls^13^. Furthermore, correct and error trials differ in their level of delay period activity. Our current results show that the activity of FS neurons that generate persistent discharges is also predictive of behavior, particularly in the prefrontal cortex, no less than of RS neurons. Classification of cells into putative pyramidal neurons and interneurons is not perfectly precise and this determination cannot be made perfectly accurately for any single neuron, however neurons classified as FS are more likely to correspond to interneurons, allowing for meaningful comparisons between populations ^17^.

### Roles of FS and RS neurons in cognitive functions

Fast Spiking and Regular Spiking neurons are activated in a range of cognitive functions, however their roles are often divergent. Prefrontal Regular Spiking neurons (also referred to as Broad-Spiking in the literature) generally exhibit smaller receptive fields than Fast Spiking neurons^17, 31^. RS and FS neurons are also differentially activated in attention tasks, with FS neurons much more sensitive to task^16, 32^. Finally, FS and RS neurons are differentially affected by training to perform a new cognitive task, with FS firing rate elicited by stimulus presentations increasing to a greater extent^3, 4^. Our current results add to the list of tasks where FS neurons have been shown to exhibit persistent activity. Importantly, we show that both FS and RS neurons that generate persistent discharges are predictive of behavior in these tasks, consistent with predictions of the bump attractor model, and more broadly with the idea that persistent activity determines behavior in working memory tasks.

### Prefrontal and parietal roles in working memory

Numerous studies have implicated the prefrontal cortex as being critical for the maintenance of working memory^33^. However, evidence has accumulated of other brain areas playing a role in working memory, including the PPC, to the extent that is commonly referred to as part of the fronto-parietal network^34^. Similar patterns of activation have been observed across PFC and PPC areas during working memory tasks^35^. Studies have thus emphasized that cognitive performance relies heavily on a robust, distributed neural network across both areas^36^.

Prefrontal and parietal neurons do exhibit some specialization. PPC neurons are generally less able to resist the effect of distracting stimuli^37, 38^, though the specific patterns of responses in the two areas depend on the exact task a subject is performing^14, 39^. Persistent activity in prefrontal cortex also appears more robust and less variable from trial to trial^40–42^. Such differences in functional properties can be traced to differences in intrinsic properties of neurons and circuits in the two areas, including the relative frequency of different interneuron types in the prefrontal cortex^43–46^. Our results also raise the possibility that the activity of FS neurons in the prefrontal cortex is more reliably related to subject behavior, though our sample size was small to make this determination definitively; future studies may further test this idea.

